# Detection of outlier methylation from bisulfite sequencing data with novel Bioconductor package BOREALIS

**DOI:** 10.1101/2022.05.19.492700

**Authors:** Gavin R. Oliver, Garrett Jenkinson, Rory J. Olson, Laura E. Schultz-Rogers, Eric W. Klee

**Affiliations:** Department of Quantitative Health Sciences, Mayo Clinic, Rochester, MN, USA; Center for Individualized Medicine, Mayo Clinic, Rochester, MN, USA

**Keywords:** DNA methylation, bisulfite sequencing, outlier analysis, rare disease

## Abstract

DNA sequencing results in genetic diagnosis of 18-40% of previously unsolved cases, while the incorporation of RNA-Seq analysis has more recently been shown to generate significant numbers of previously unattainable diagnoses. Multiple inborn diseases resulting from disorders of genomic imprinting are well characterized and a growing body of literature suggest the causative or correlative role of aberrant DNA methylation in diverse rare inherited conditions. Therefore, the systematic application of genomic-wide methylation-based sequencing for undiagnosed cases of rare disease is a logical progression from current testing paradigms. Following the rationale previously exploited in RNA-based studies of rare disease, we can assume that disease-associated methylation aberrations in an individual will demonstrate significant differences from individuals with unrelated phenotypes. Thus, aberrantly methylated sites will be outliers from a heterogeneous cohort of individuals.

Based on this rationale, we present BOREALIS: **B**isulfite-seq **O**utlie**R** M**E**thylation **A**t Sing**L**e-S**I**te Re**S**olution. BOREALIS uses a beta binomial model to identify outlier methylation at single CpG site resolution from bisulfite sequencing data. This method addresses a need unmet by standard differential methylation analyses based on case-control groups. Utilizing a heterogeneous cohort of 94 rare disease patients undiagnosed following DNA-based testing we show that BOREALIS can successfully identify outlier methylation linked to phenotypically relevant genes, providing a new avenue of exploration in the quest for increased diagnostic rates in rare disease patients. We highlight the case of a patient with previously undetected hypermethylation patterns that are informing clinical decision-making. BOREALIS is implemented in R and is freely available as a Bioconductor package.

## 1 Introduction

Human DNA methylation is a mitotically heritable epigenetic mark involving the covalent addition of a methyl group to the carbon-5 position of cytosine molecules occurring adjacent to and upstream of guanine molecules (CpG sites). DNA methylation plays well-documented roles in embryonic development, genomic imprinting, X-chromosome inactivation and retrotransposon silencing(1, 2). Functionally, DNA methylation patterning has been shown to affect chromatin plasticity, gene expression, and more recently, patterns of exon inclusion in transcribed RNA(3, 4). DNA methylation profiles have been demonstrated to track biological age(5–8) and have been extensively studied in pan-cancer analyses(9, 10), suggesting a functional role in oncogenesis. Multiple inborn diseases resulting from disorders of genomic imprinting are well characterized, such as the Prader-Willi or Angelman Syndrome (11). Furthermore, a growing body of literature also suggests associations between DNA methylation and conditions including Parkinson’s Disease(12), Alzheimer’s(5, 13), Irritable Bowel Disease(14), autism(15), chronic pain(16), obsessive compulsive disorder(17), depression(18), suicidality (19) and more.

Early characterization of DNA methylation patterns focused on wide regions of the genome, particularly regions of high CpG-site concentration (CpG islands) that are often associated with regulatory elements such as gene promoters or enhancers (3). A frequent observation in healthy tissues around transcriptionally active genes is a methylation landscape that has little methylation in or around promoters, and more methylation in gene body exons and intergenic regions; conversely, transcriptionally inactive genes tend to show increased methylation within their promoter region and less in the gene body. In cancer for example, it has been shown that methylation patterns show extensive deviation from those normally observed (1). This leads to globally hypomethylated regions and accompanying increases in DNA methylation stochasticity (20, 21). These alterations are often associated with multiple mutations in epigenetic maintenance enzymes (22) although extensive methylation aberrations are observed even in cancers with low mutational burden (20, 23, 24). While microarray-based methylation profiling continues to be utilized with success, the advent of next-generation sequencing has enabled genome-wide methylation analysis at single-site resolution and at greatly increased number of genomic CpG sites (25). This has enabled more complex pictures of methylation patterning to emerge, including the association of multiple specific-site or even single-site methylation with developmental processes (26) or disease (27–35). It is likely that high-resolution methylation profiling will continue to reveal the functional importance of nuanced focal methylation alterations, as well as differential patterning across broader genomic features.

Like the study of methylation, investigations of rare genetic disease have benefited from the era of high throughput sequencing. DNA sequencing alone has been reported to result in genetic diagnosis of 18-40% of cases previously unsolved using standard clinical testing (36–38). Recent years have seen concerted efforts to bolster rates of diagnosis through incorporation of RNA-Seq analysis into the diagnostic toolkit. A variety of testing paradigms have been investigated and shown measurable benefit in supplementing DNA testing. RNA fusion transcription, allele-specific expression, alternative splicing analysis and gene expression quantification have each been applied and generated diagnoses previously unattainable with DNA alone (39–42).

With the growing body of evidence linking methylation to disease, the systematic application of genome-wide methylation-based sequencing for undiagnosed cases of suspected rare inherited disease is a logical progression from current DNA and RNA testing paradigms (43, 44). Array-based ‘epi-signature’ testing is already clinically available for some 40 inherited conditions affecting around 60 genes (45, 46). Methylation-based studies have also suggested the causative or correlative role of aberrant methylation in diverse rare inherited conditions including Kabuki Syndrome (47), Werner Syndrome(48) rare ophthalmic disease (49), orofacial cleft (50) and fucosidosis (51), either in the presence or absence of known DNA mutations. Whether deviation from a normal methylation pattern reflects alterations of genetic or epigenetic origin, expanded methylation profiling could offer the ability to detect diagnostic signals unique to the epigenome or undetectable in DNA and RNA due to lack of measurable manifestation in those materials, or due to shortcomings in current technologies or analytical approaches.

Since functionally relevant DNA methylation aberrations might occur at a single-site, multiple specific-site or regional scale, a method to detect deviant methylation should offer the ability to profile at the single-site level while enabling flexibility to consolidate calls across regions defined in the context of an individual analysis. Furthermore, in the context of rare inherited disease we can assume that disease-associated or causative methylation aberrations in a single individual will be demonstrate significant differences from other individuals with unrelated phenotypes i.e., that the aberrantly methylated sites will be outliers from a heterogeneous cohort of individuals. Similar rationale has been successfully employed in outlier expression-based RNA analysis (52–54). However, existing solutions for detection of differentially methylated CpG sites from bisulfite sequencing focus on traditional group vs group or multi-group experimental designs (55–57) and are therefore not suited to rare disease or other outlier-based analyses.

Here we present BOREALIS: **B**isulfite-seq **O**utlie**R** M**E**thylation **A**t Sing**L**e-S**I**te Re**S**olution. BOREALIS uses a beta-binomial model to identify outlier methylation at single CpG site resolution from bisulfite sequencing data, by comparing a single individual to a cohort. The method is suited for single or multi-site downstream analysis. Utilizing synthetic data as well as a heterogeneous cohort of 94 rare disease patients undiagnosed following DNA-based testing we show that BOREALIS can successfully identify outlier methylation with high statistical power, providing a new avenue of exploration in the quest for increased diagnostic rates in rare disease patients. BOREALIS is implemented in R and is freely available as a Bioconductor package.

## Materials and Methods

### BOREALIS statistical model

At a given CpG site, we assume that we have data in the form of methylated counts x_i_ and total read counts n_i_ for individuals i=1,…,I in our cohort of size I. If every individual in the population had the exact same probability p of methylation at this site, i.e., p_1_ = p_2_ = ⋯ p_I_ = p where pi is the (true-but-unknown) probability of methylation for the i^th^ individual at this site, then the methylated counts xi would be binomially distributed with parameters p and n_i_. However, we expect varying degrees of sample-to-sample variability in the probability of methylation at a given site even in a healthy cohort. Therefore, it would be more biologically accurate to assume that p_i_ for i=1,…,I have been sampled from a distribution over the unit interval. A common and mathematically convenient choice for this distribution is a beta distribution with parameters α and β. Thus the observed number of methylated reads x_i_ for the i^th^ individual can be viewed as being generated from a two-step process whereby the probability of methylation is selected p_i_ ~ Beta(α,β) and given this probability the number of methylated reads is binomially distributed x_i_|p_i_ ~ Binomial(p_i_,n_i_). Viewed in this way, we say the methylated reads are beta-binomially distributed x_i_ ~ Beta-Binomial(n_i_,α,β) with parameters α and β, and popular packages for differential DNA-methylation detection such as DSS (58) use this same distribution for methylated counts. What BOREALIS does differently from traditional tools such as DSS (57) is that it builds its statistical model explicitly for the purpose of outlier detection compared to a cohort, which requires alternative statistical framing and considerations as compared to group-versus-group analyses (53) as illustrated in Figure 1. Specifically, at each CpG site, BOREALIS takes the input data {(x_i_,n_i_): i=1,…,I} and estimates the population-level parameters α and β using the R package gamlss (58). In practice, we implement Laplace Smoothing on the counts (i.e., we use as counts 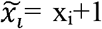 and 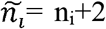) as a regularization step to help deal with any samples with low counts. Then from these estimated α and β parameters, we can for the i^th^ sample compute the left-sided p-value by looking at the probability that a value of x_i_ or fewer methylated reads were generated from a Beta-Binomial(n_i_,α,β), and likewise a right-sided p-value would evaluate the probability that a value of x_i_ or greater methylated reads came from this distribution. We implement this probability calculation using the pBB function of the gamlss package in R. The two-sided p-value is computed as two times the lesser of these one-sided p-values.

**Figure 1.**
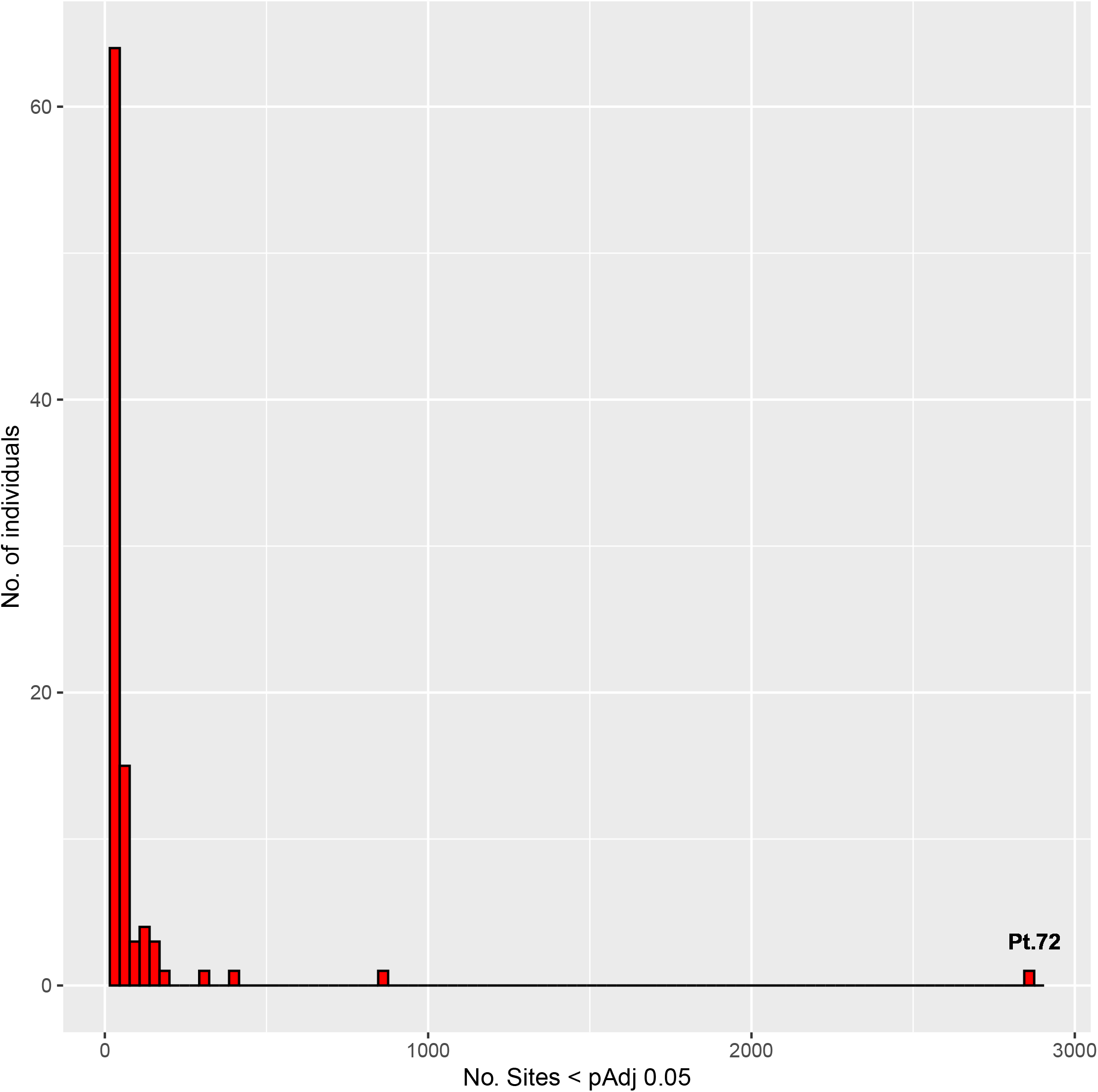
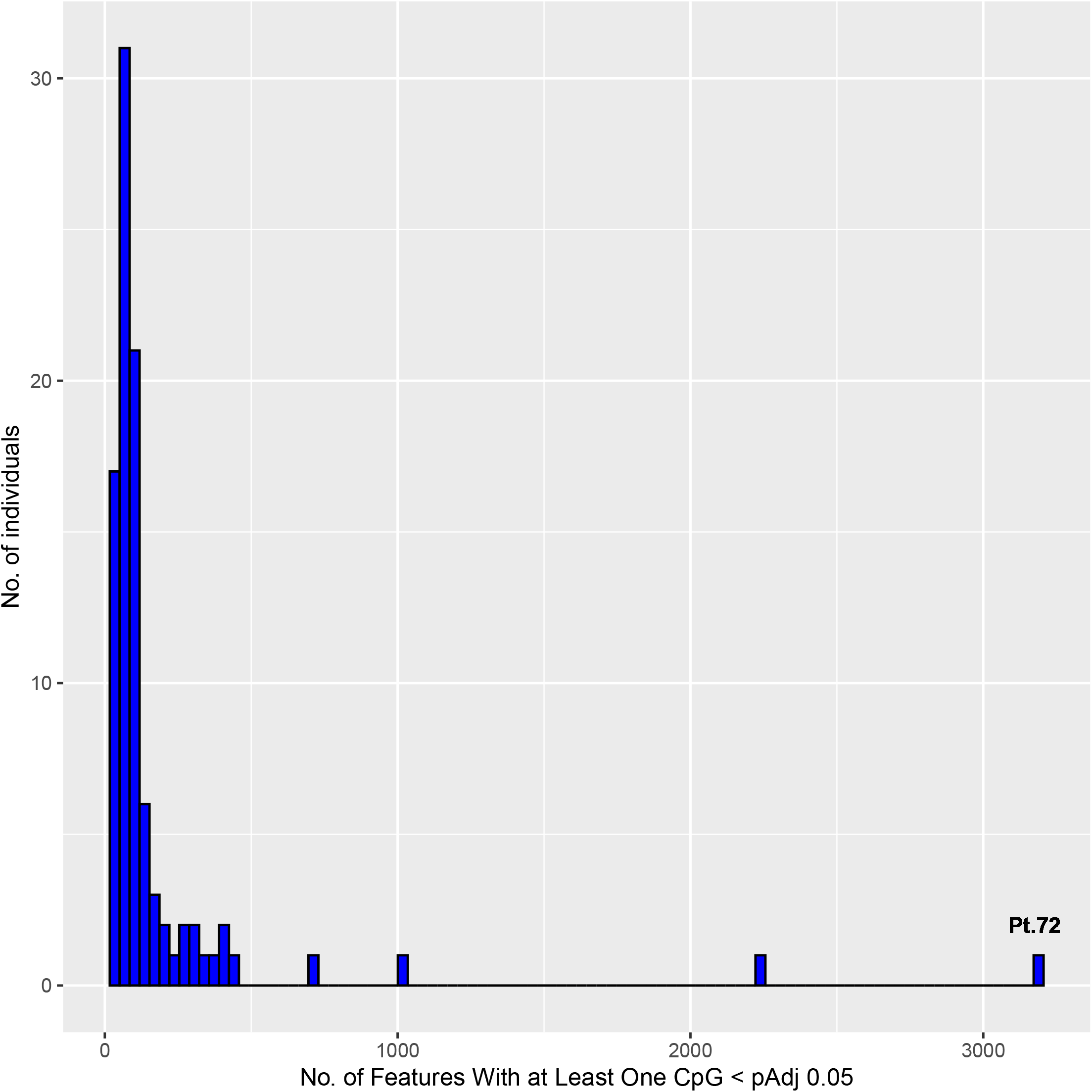
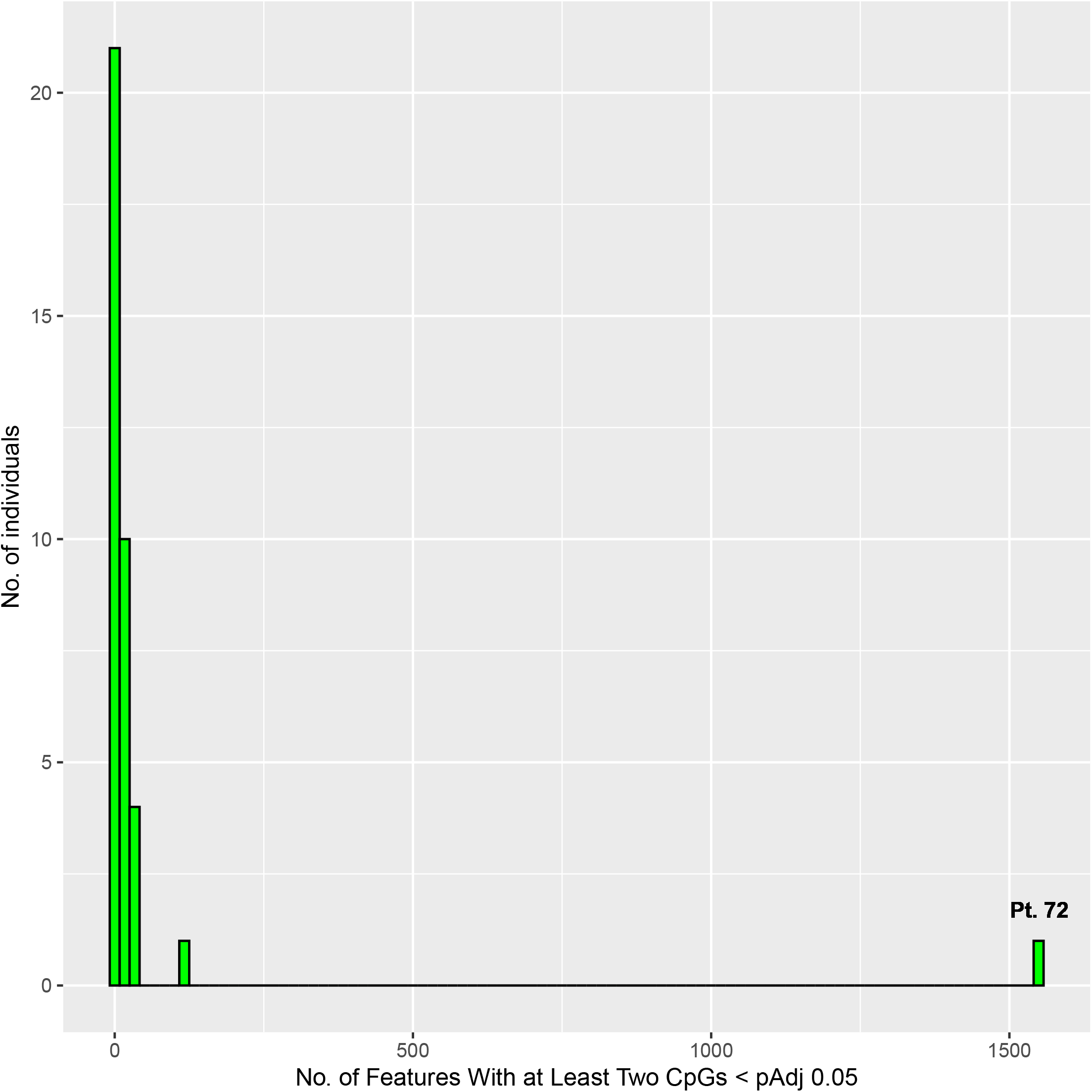
Conceptual differences between traditional case vs control analysis and the BOREALIS approach. While traditional approaches to differential methylation analysis are based on group case vs. control analysis, BOREALIS utilizes a one vs. many outlier approach whereby individual(s) are compared to a cohort. This approach is especially useful when multiple similar cases are difficult to identify, as is the case in rare disease studies. By comparing every affected individual to a cohort of heterogeneous individuals, outlier methylation can be identified at individual CpG sites for all members of the cohort, without the requirement for multiple similar cases.

### Monte Carlo power simulations

We performed Monte Carlo simulations of an outlier sample in cohorts of varying sizes sequenced at varying depths of coverage. Namely, after selecting a cohort size *I* and an average depth of coverage *D* we conducted 10,000 Monte Carlo simulations wherein each sample *i* in the cohort has sequencing depth *d_i_* drawn from a Poisson distribution with mean *D*. The number of methylated reads in cohort sample *i* is drawn from a Beta-Binomial distribution with mean 0.8 and dispersion 0.1, and then these simulated cohort data are fit using the BOREALIS model. An outlier sample is then simulated with sequencing depth *d* drawn from a Poisson distribution with mean *D* and number of methylated reads given by a Binomial distribution with mean 0.3. BOREALIS is then used to compute a p-value thresholded at level 0.05, and the power is given by the fraction of the 10,000 simulations that correctly reject the null hypothesis.

### Cohort selection

A total of 94 individuals seeking diagnosis of suspected inherited disease were selected, having undergone genetic consultation at Mayo Clinic and referral to the Center for Individualized Medicine. All individuals had previously undergone genomic sequencing without successful genetic diagnosis.

### Ethical compliance

Study subjects and families provided written informed consent to a research protocol approved by the Mayo Clinic Institutional Review Board for this study.

### Methylation profiling

Reduced representation bisulfite sequencing (RRBS) was performed on peripheral blood mononuclear cells collected from all 94 individuals. Samples were first randomized for sex, age and phenotype. RRBS libraries were then prepared with 100 ng of genomic DNA using the NuGen RRBS Ovation Kit (NuGen, Redwood City, CA). Briefly, dsDNA was digested with Msp1 and indexed methylated adaptors were ligated to the digested fragments with T4 DNA ligase. Ligated DNA was repaired with Final Repair mix. Bisulfite modification was performed using the EZ-DNA Methylation Kit (Zymo Research, Irvine, CA). Bisulfite modified product was then amplified with PCR and purified with AMPure beads. The concentration and size distribution of the completed libraries were determined using the Fragment Analyzer™ Standard Sensitivity NGS Fragment Analysis kit (Agilent, Santa Clara, CA) and Qubit fluorometry (Invitrogen, Carlsbad, CA). Completed libraries were pooled and 20% commercially prepared PhiX library (Illumina) was added to increase base diversity for improved sequencing quality. Samples were sequenced at eight samples per lane, following Illumina’s standard protocol using the Illumina cBot and HiSeq 3000/4000 PE Cluster Kit. A custom Read 1 primer, MetSeq Read 1 (original concentration 100uM), was spiked in with the Illumina Read 1 primer. The flow cells were sequenced as 51bp paired end reads on an Illumina HiSeq 4000 using the HiSeq 3000/4000 sequencing kit and HCS v3.4.038 collection software. Base-calling was performed using Illumina’s RTA version 2.7.7.

### RRBS data processing and analysis

RRBS reads were aligned to the human genome (hg19) using Bismark v0.22.3 (59). Bisulfite conversion efficiency was estimated in each sample as 100%-X where X was the global precent methylation in CHH context cytosines. Samples are considered to pass quality control if bisulfite conversion efficiency was greater than 99% Reads were inspected using Bismark M-bias plots and problematic end regions displaying methylation bias were hard-trimmed using Trimgalore v0.6.5 before realignment. Trimmed and aligned data were subsequently processed with BOREALIS, producing single-site resolution outlier methylation data for each of the 94 samples. CpG sites covered by fewer than 10 reads in a given sample were excluded from result generation in that sample. CpG sites were annotated with their corresponding genomic features using the Bioconductor package annotatr (60). Chromosomal enrichment testing for affected genes was performed using Enrichr (63). P-values were computed from a Fisher exact test. Other data summarization and exploration were conducted in R.

### Phenotype-based gene prioritization

We applied an *in-silico* phenotype-based gene prioritization method named PCAN (Phenotype consensus analysis to support disease-gene association) (61) to annotate confidence in known links between a gene and a patient’s phenotype. PCAN uses semantic similarity scoring to measure closeness between the phenotypic terms mutually associated with a patient and a gene. Scores are ranked by simultaneously measuring semantic similarity for all disease-associated genes in the ClinVar database (62) versus each patient’s phenotype and producing a rank (e.g. 10 indicates that a gene produces a semantic similarity score ranked 10^th^ when compared to all other disease-linked ClinVar genes). This method enables rapid prioritization of results from high-throughput analysis without the need for extensive manual curation of genes.

### RNA Sequencing

Sequencing was conducted on whole blood. Blood-derived RNA was obtained by collecting peripheral whole blood in PAXgene blood RNA tubes and using the QIAcube system (Qiagen) according to the manufacturer’s protocol for RNA extraction. Sequencing libraries were prepared with the TruSeq RNA Access Library Prep Kit (Illumina, San Diego, CA). Paired-end 101-basepair reads were sequenced on an Illumina HiSeq 2500 using the TruSeq Rapid SBS sequencing kit version 1 and HCS version 2.0.12.0 data collection software. Base calling was performed using Illumina’s RTA version 1.17.21.3.

### Outlier RNA expression analysis

RNA-sequencing analysis was performed using MAP-RSeq (64). Reads were aligned to the human genome (hg19) and transcriptome using Tophat2(65) running Bowtie (v1)(66). Gene level raw read counts and RPKMs were generated using HiSeq(67) and BedTools (68), respectively. Genes whose 0.95 quantile RPKM was less than 1 or whose counts were zero in more than a quarter of samples were excluded from the analysis. Outlier expression analysis was then performed using OUTRIDER (v1.1.1)(69) with Benjamini-Hochberg correction for multiple comparisons, which provided confounder-corrected z-scores, log2 fold changes, p-values and adjusted p-values.

## Results

### Simulated data and power analysis

To evaluate the statistical power of the proposed approach under a variety of conditions, we performed Monte Carlo simulations of an outlier sample in a cohort of varying sizes sequenced at varying depths of coverage. As described in the Methods, we conducted 10,000 Monte Carlo simulations for each sequencing depth and cohort size. In these simulations, we randomly sampled the sequencing depth of each sample, as well as the proportion or reads that were methylated. The BOREALIS algorithm was then used to build its beta-binomial model for each simulated cohort and an outlier sample is then simulated with BOREALIS used to compute a p-value. The power is estimated by the fraction of the 10,000 simulations that correctly reject the null hypothesis at a level 0.05. The results of this procedure are plotted in Figure 2A, demonstrating that BOREALIS can accurately detect outlier methylation events in modest cohort sizes and sequencing depths. To provide users with the ability to visually review the methylation distributions underlying any call made by BOREALIS, we provide a built-in plotting function whose outputs are illustrated in Figure 2B.

**Figure 2.**
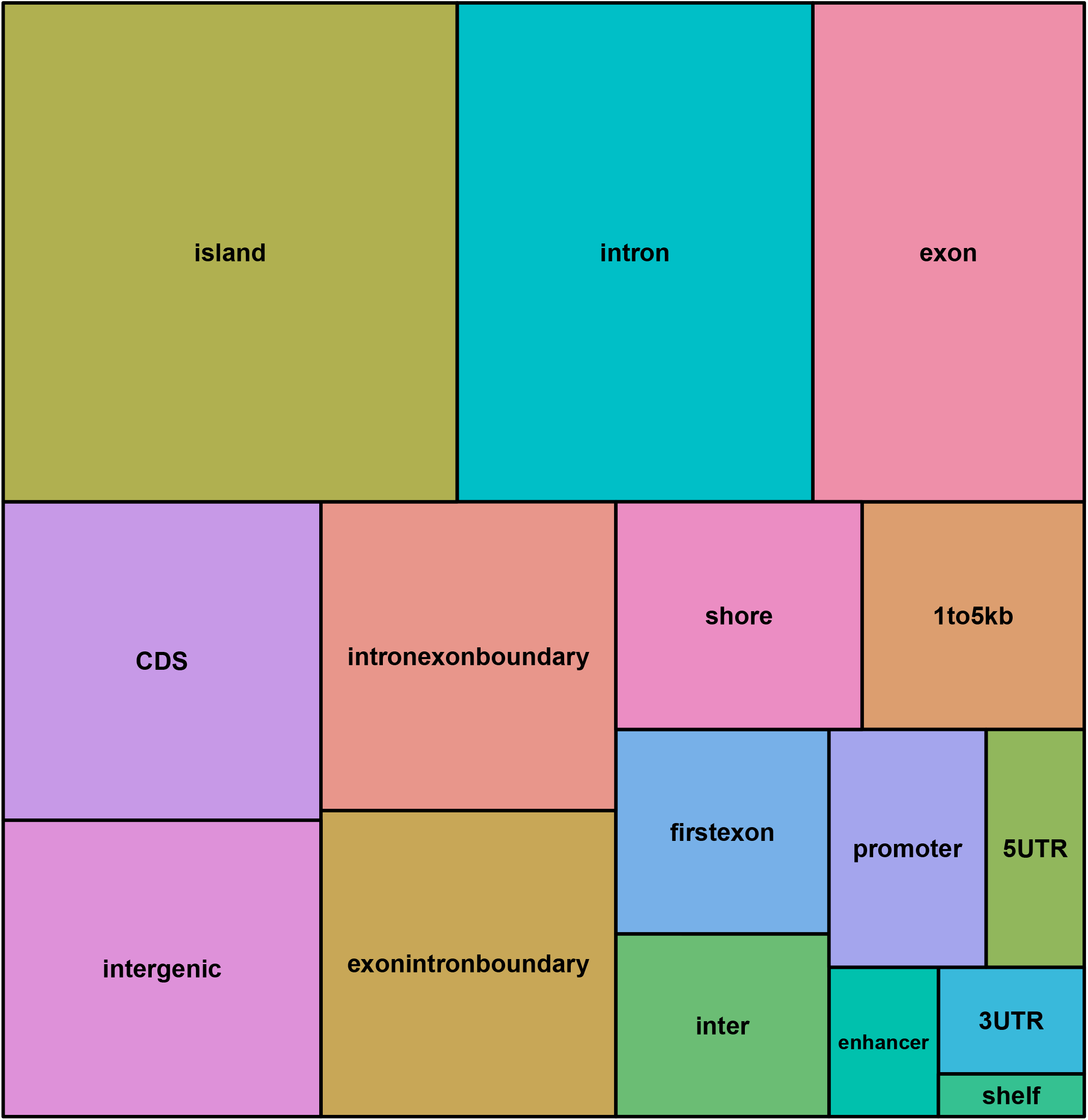
BOREALIS power analysis and single site methylation profile output by BOREALIS. A) Graphical summarization of Monte Carlo simulations of an outlier sample in a cohort of varying sizes and depths of sequencing coverage. Ten thousand simulations were performed for each set of experimental conditions whereby random sampling of sequencing depth for each sample and the proportion methylated reads was performed. BOREALIS built its beta-binomial model for each simulated cohort and an outlier sample was simulated with BOREALIS used to compute a p-value. Power estimation is based on the simulations correctly rejecting the null hypothesis at a level p<=0.05. The parameters mu (mean methylation fraction at a given site), sigma (variability in methylation fraction at a given site) and muAb (deviation from the mean methylation level in the outlier sample) are fixed for the purposes of the simulation shown. B) The BOREALIS package includes functionality to visually represent the methylation profile at any given CpG site, as compared to the cohort. Here a single site within the *LTB4R* gene promoter is shown for Patient 72.

### Rare disease cohort

The patient cohort consisted of 56 males and 38 females. Ages at initial referral ranged from 0-19 years with a median age of 8.5 years. Clinical presentations were heterogeneous and comprised a spectrum of neurological, muscular, gastrointestinal, immune, skeletal, and connective tissue disorders.

### Read mapping and quality control

Full read mapping and QC information are provided in Supplementary Table 1. An average of approximately 30 million read pairs (+/− 1.2 million) were generated per sample. Consistent mapping efficiencies were consistently observed for all samples. An average of 338 million Cytosines (+/− 16.2 million) were analyzed per sample. Low levels of Cytosine methylation in CHH context (maximum 0.5%) were observed across all samples, confirming adequate and consistent bisulfite conversion efficiencies of at least 99.5%.

### Cohort methylation summary statistics

A total of 1.93 million unique CpG sites were profiled for outlier methylation by BOREALIS following the processing described in Methods. Sequencing depth across samples showed a median of 21 reads per CpG site (+/− 12 reads). The median mu value (average methylation level per-CpG site, across all samples) was 62.6% (+/− 42.1%). The theta value utilized as a measurement of per-CpG site methylation variability across samples had a median of 0 (+/− 0), demonstrating predominantly close regulation of per CpG site methylation across the cohort. Individual CpG site methylation levels ranged from 0 to 100% with a median methylation level of 12% (+/− 17.8%). Effect sizes at each CpG site per-patient versus the cohort, varied from 0 - 92.9% (Median = 5.3% +/− 3.8%)

### Number of outlier CpG sites per sample

We utilized a Benjamini-Hochberg corrected p-value threshold of <= 0.05 to define significant outlier CpG sites within the cohort. The number of significant CpGs ranged from 16 – 2,858 per case (Median = 36 +/− 14). The distribution of the number of significant sites per patient are shown in Figure 3. Since cases with low numbers of significant sites are most similar to the cohort, and therefore least likely to be clinically relevant, we focused on those with unusually high numbers of significant outlier CpG sites. Patient 72 was particularly notable as a distant outlier with 2,858 significant sites versus 869 and 399 sites for its nearest neighbors, Patient 36 and Patient 55 respectively.

**Figure 3.**
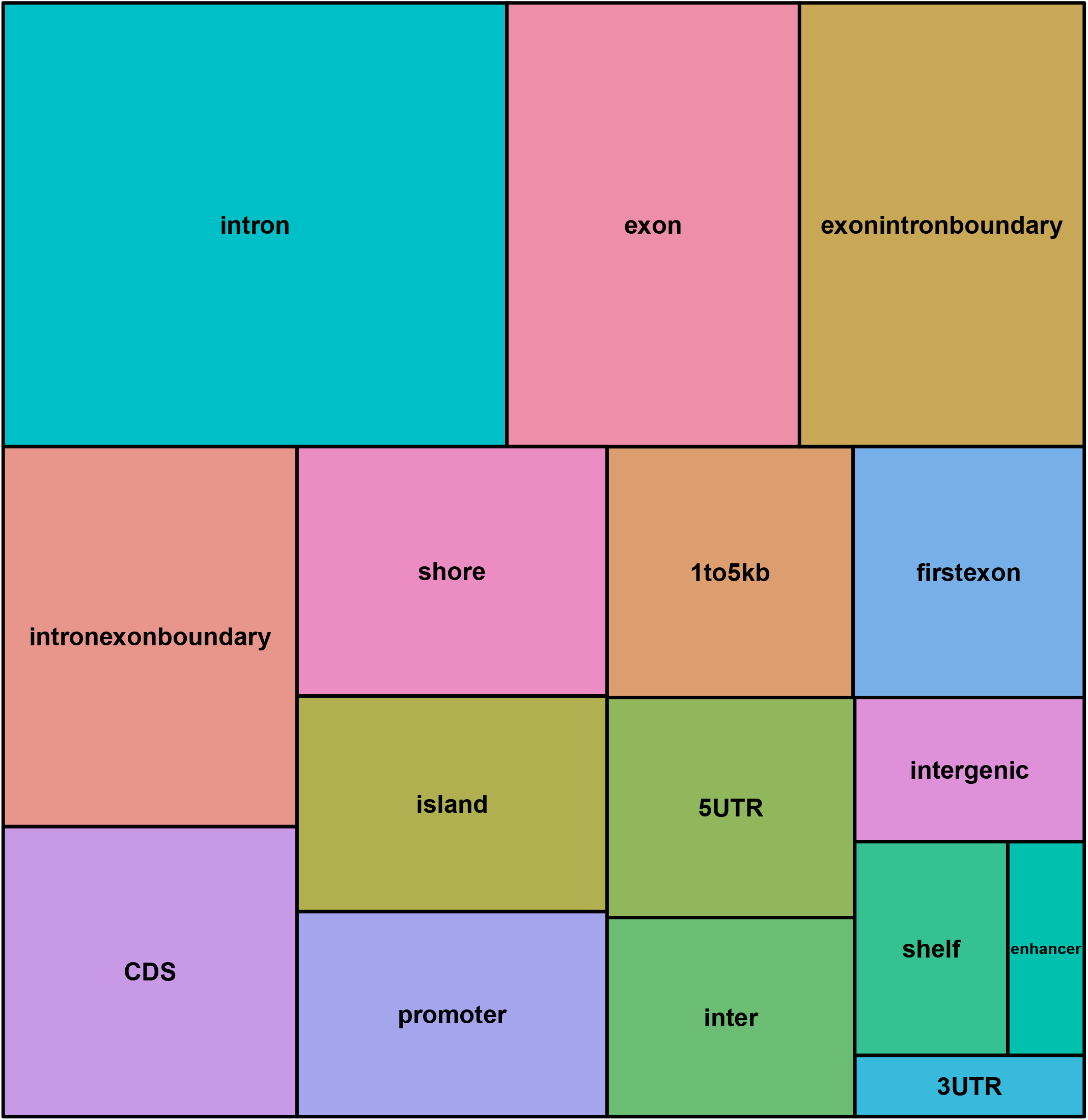
Number of significant outlier CpG sites per case. Histogram of the number of significant outlier CpG sites per case. Patient 72 is a notable outlier with many more outlier CpG sites overall compared to other patients.

To determine if outlier samples appeared affected genome-wide, we performed a per-chromosome summarization of outlier sites. Patient 72 was the most significant outlier based on number of significant outlier CpG sites across 22 chromosomes. Patients 36 and 55 were 2^nd^ or 3^rd^ most significant outlier across 22 and 11 chromosomes respectively, indicating potentially widespread or global methylation aberrations for each outlier sample. Eleven other patients appeared as top three outliers for 1-4 chromosomes per patient.

### Number of features per case with outlier CpG sites

A median of 410,034 (+/− 14,180) unique epigenetically relevant features per case were represented in the data output by BOREALIS. These consisted of common genomic features with known links to epigenetic regulation and included CpG Islands, Promoters, 5′ and 3′ UTRs, enhancers, CpG Island Shores and more. The number of unique gene promoters annotated per case ranged from 19,432 – 22,888 (Median 21,638 +/− 524) All annotated feature classes and their relative proportions are visually summarized in Supplementary Figure 1.

We calculated the number of annotated genomic features per sample with (a) one or more significant outlier sites per feature and (b) two or more significant outlier sites per feature. For analysis (a) the number of affected features per patient ranged from 31 to 3186 (Median=79.5 +/− 43), affecting 8 to 744 genes per patient. Patients 72 was again a notable outlier with 3,186 affected features versus 2,250 and 1,019 features for its nearest neighbors, Patient 36 and 55 respectively. For analysis (b), the number of affected features per patient ranged from 1 to 1,547 (Median=7 +/− 4.45), affecting 1 to 298 genes per patient. Patient 72 (1,547 features) was again a notable outlier. The distributions for each analysis are shown in Figure 4 and 5.

**Figure 4.**
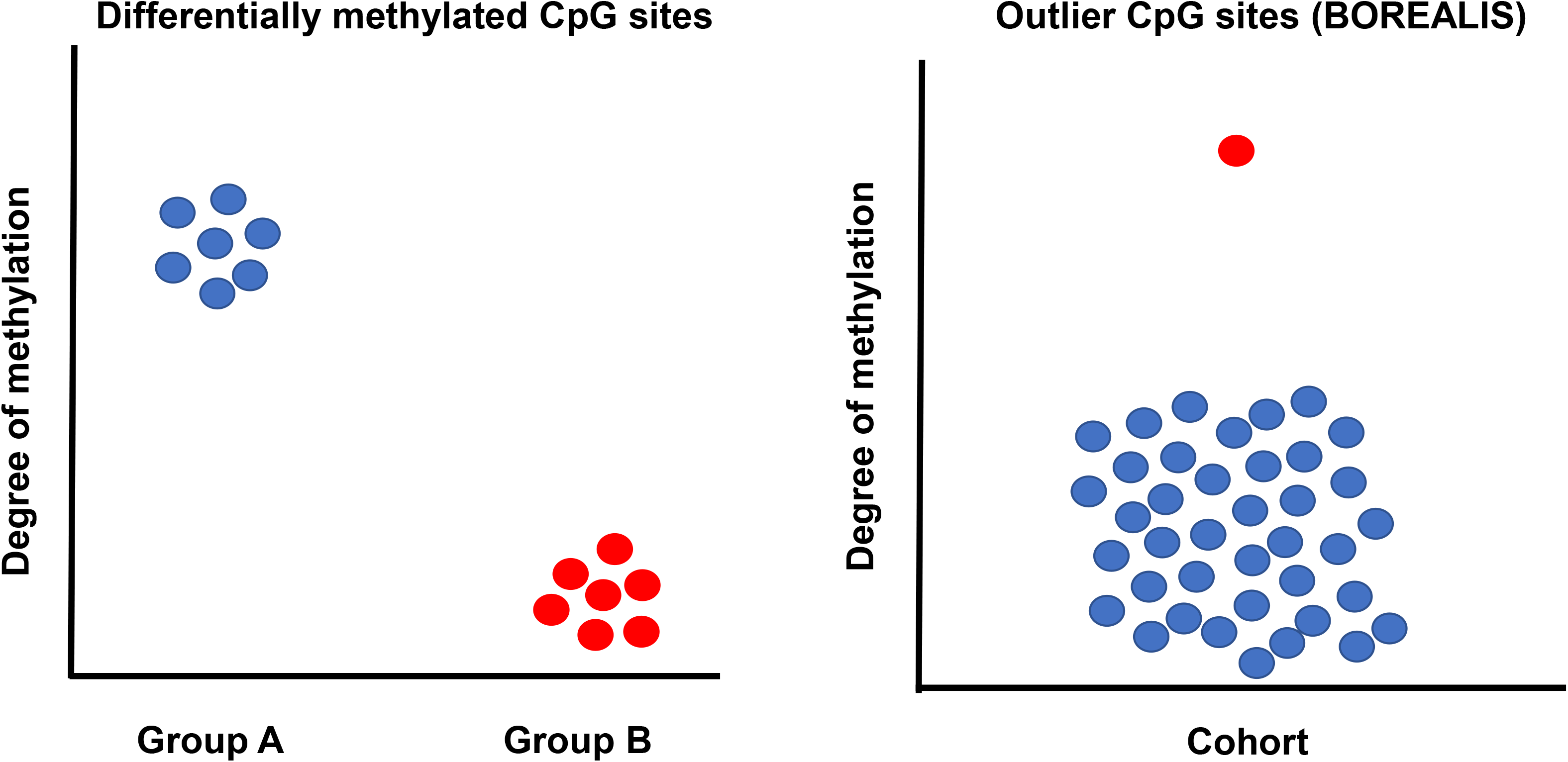
Histogram of the number of features containing one or more significant CpG sites. Patient 72 is a notable outlier with many more affected features than other patients.

**Figure 5.**
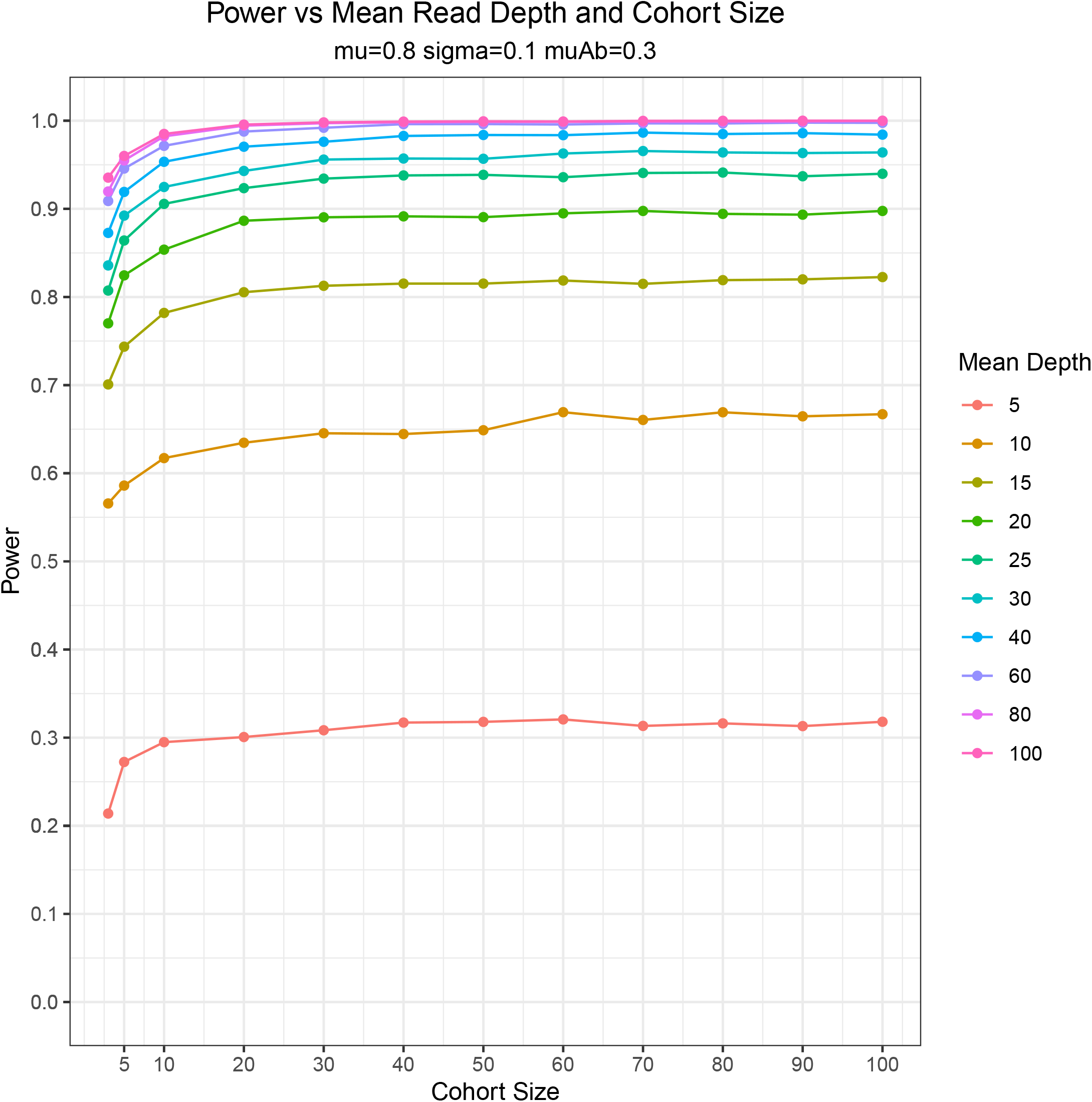
Histogram of the number of features containing one or more significant CpG sites. Patient 72 is a notable outlier with many more affected features than other patients.

### Number of significant sites per epigenetic feature

We wished to further characterize the number of significant outlier sites within epigenetic features. The number of significant outlier CpG sites per annotated feature ranged from 0 - 36 across the cohort (Median = 0 +/− 0). A total of 24,949 annotated features overlapped at least one significant outlier CpG site. The majority (16,979 features) had only one significant CpG site while 2,019 had two significant sites. A total of 1499 features had ten or more significant sites while only 30 overlapped thirty or more significant sites. A full breakdown of the number of significant outlier sites per feature across the cohort is provided in Supplementary Table 2.

Notably, features containing larger numbers of significant outlier sites skewed heavily toward Patient 72. Only Patient 72 possessed features with greater than 16 significant sites per feature. Although 15 patients possessed features with five or more significant sites, Patient 72 had 615 such features, while the other patients had only 3 – 15 (Median=6 +/− 2.22). A full summary of number of features per patient with varying number of significant sites is provided in Supplementary Table 3.

### Hypermethylation vs hypomethylation

We wished to determine the percentage of significant outlier sites per patient called hypermethylated vs hypomethylated by BOREALIS. Hypermethylation percentages for the cohort ranged from 0-96.5% (Median=3.92% +/− 3.93%). Interestingly, Patient 72 was a distant outlier compared to all other samples with its hypermethylation percentage of 96.5% falling well beyond its closest neighbors Patient 54 (41.3%) and Patient 91 (41.2%). A total of 30 samples demonstrated a 0% hypermethylation percentage in their significant outlier calls. Hypermethylation percentages for the cohort are include in Supplementary Table 4.

Of the 2,019 features containing more than one significant outlier CpG site in the cohort, almost all (1,995) showed concordance in directionality of sites affected i.e. all were either hypermethylated or hypomethylated compared to the cohort. Only 24 features contained outlier CpG sites with discordant directionality and 18 of these contained only two outlier sites (i.e. one hypermethylated and one hypomethylated). The remaining six features consisted of one intergenic region in Patient 36, as well as three overlapping *GLI2* introns, and two distinct intergenic regions in Patient 72. The number of significant outlier CpGs in these regions ranged from 4 – 13 in number, with 1 – 2 discordant CpGs per feature.

### Patient 72 case study

We selected Patient 72 to perform further follow-up investigation based on their status as an outlier in terms of total number of significant sites, % hypermethylated sites and total number of genomic features affected.

Patient 72 is a male child who presented to Mayo Clinic’s Center for Individualized medicine at 8 years of age, suffering from a broad phenotypic history including neonatal hypotonia, ptosis, poor suck, obsessive-compulsive behavior, plagiocephaly, global developmental delay, ectopic kidney, astigmatism, behavioral abnormality and dysplasia of the second lumbar vertebra. The patient had previously undergone exome sequencing but no candidate DNA variants with links to the phenotype were identified.

We wished to determine if the aberrant methylation profile specifically affected any genes with established ties to the patient’s phenotype. Patient 72 had 1,547 such features with more than a single outlier CpG site, and these features were distributed across 298 genes. A graphical summary of the affected feature classes is provided in Figure 6.

**Figure 6.**
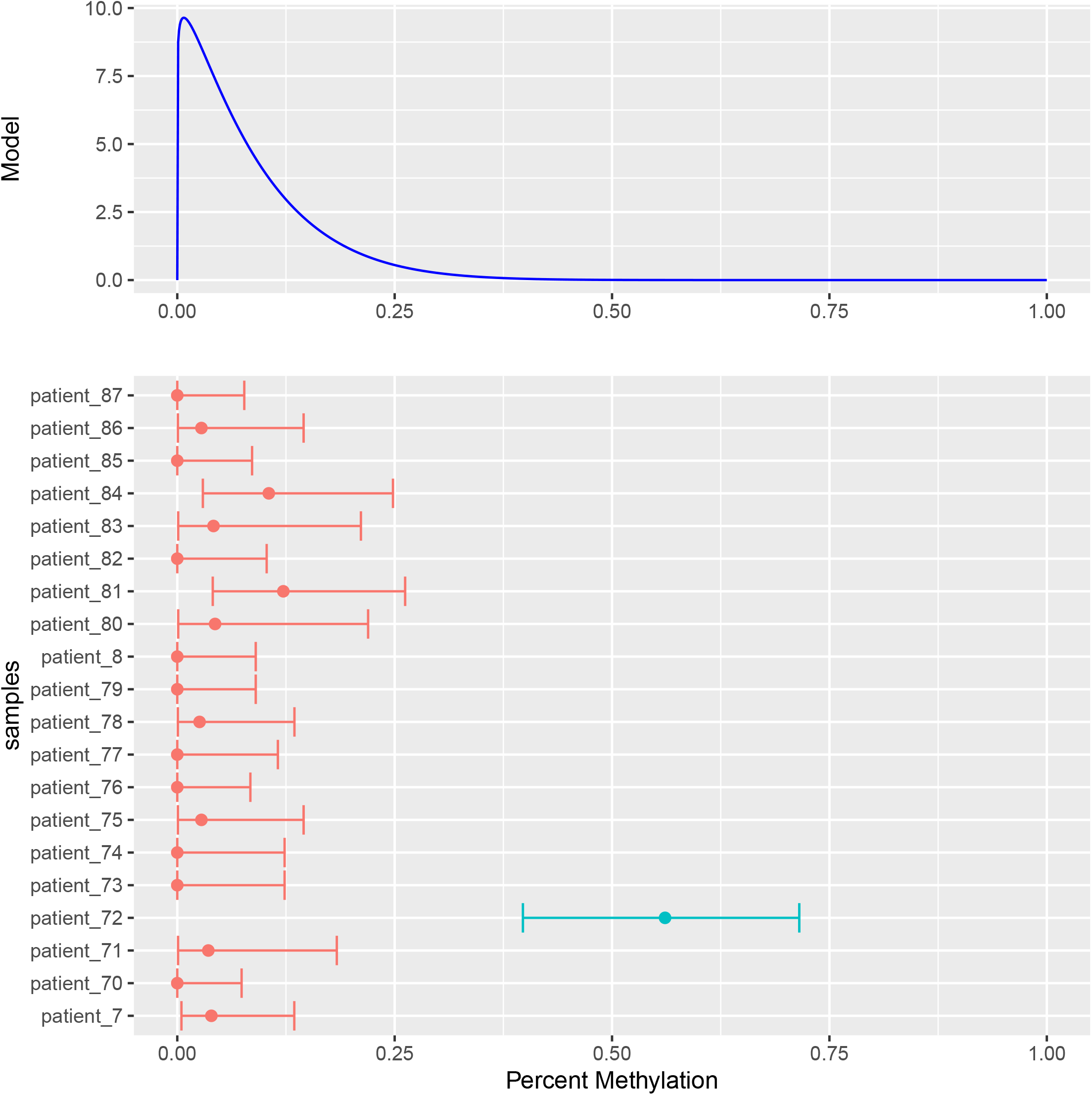
Treemap showing the relative proportions of different genomic features containing two or more significant outlier CpG sites in Patient 72. Features were annotated using the Bioconductor package “annotatr” as described in the the BOREALIS Bioconductor vignette. A total of 1,547 such features distributed were across 298 genes.

We applied phenotype-based semantic similarity scoring to all 298 genes with PCAN to provide a measure of how closely the phenotype known to be associated with each gene matches the patient’s phenotype. Possible PCAN ranks range from 1 (very strongly linked) to 3737 (links essentially non-existent). Genes with no known clinical phenotypic evidence are not scored.

PCAN ranks for the 298 genes ranged from 44 – 3737. Using a cutoff of rank 100, only two genes were included. These were *GLI2* (rank 97) and *PCSK9* (rank 44). *PCSK9* overlapped two significant outlier CpGs in a region corresponding to an intron and a CpG shore, while *GLI2* contained 13 significant outlier CpGs in region corresponding to intronic sequence and a CpG island. *GLI2* is linked to Culler-Jones Syndrome and Holoprosencephaly 9, while *PCSK9* is linked to Hypercholesterolemia, familial, 3 and Low-density lipoprotein cholesterol level QTL 1. Ultimately, neither gene was considered a sufficient phenotypic match to justify further investigation of the individual results.

The feature containing the largest number of significant outlier sites was the region 1-5 kb upstream of the *KCNH2* gene (PCAN rank 353) in Patient 72. Within this feature, 163 individual CpG sites exist in the reference genome. Data was available for 53 of these, of which 36 produced significant adjusted p-values <0.05. All 36 significant CpG sites were hypermethylated (i.e. increased methylation compared to the cohort). While the result is interesting, little evidence exists to link *KCNH2* to the patient phenotype.

A total of 39 genes contained significant outlier CpGs in their promoters (Range = 2–18 CpG sites, Median = 4 +/− 2.97). None of the genes with affected promoters had evidence linking them strongly to the patient phenotype (PCAN ranks 843 – 3311).

### Outlier RNA Expression

To provide initial validation of effects on RNA transcription, we performed RNA-Seq on Patient 72 whole-blood using identical protocols to an existing internal cohort of 405 phenotypically heterogeneous rare disease RNA samples. We used the OUTRIDER package to perform outlier RNA expression analysis against this cohort. A total of 10,181 genes were sufficiently expressed in whole blood to undergo profiling. A conservative OUTRIDER analysis indicated 7 genes (*USP48, WSB1, PODXL2, CIDEB, TBX19, NLRP6* and *ZNF469*) with outlier expression (pAdj<=0.05) compared to the internal RNA cohort. Three of the seven genes (*CIDEB, NLRP6* and *ZNF469*) contained features with multiple significant outlier CpG sites, indicating a statistically significant overlap between the OUTRIDER and BOREALIS results (Fisher’s exact test p-value 2.3×10^−3^, OR=16.98). These significant outlier sites were exclusively hypermethylated and the features affected were often associated with transcriptional control, including promoters, the 4kb region upstream of promoters, 5’ UTR, first exons, and CpG islands and shores. The maximum number of significant CpG sites per feature in significant OUTRIDER genes was 18 sites for *CIDEB*, 14 sites for *ZNF469*, and 4 for *NLRP6.* All three genes appeared downregulated compared to the RNA cohort (log2 fold-change −0.92, −1.47 and −2.12 for *CIDEB, ZNF469* and *NLRP6* respectively). While efforts were made to assess the expression of other specific genes highlighted as potentially interesting by the methylation analysis (*KCNH2, GLI2, PCSK9*), analysis was not possible due to low/missing expression of these genes in whole blood.

## Discussion

We developed the Bioconductor package BOREALIS, for the detection of outlier methylation at single CpG site resolution using a single sample vs cohort experimental design, and subsequently demonstrated its utility in the context of rare disease patient cohort. To our knowledge, BOREALIS represents a first-of-its-kind computational solution for this cohort-based testing paradigm in methylation sequencing. This outlier-based experimental design has previously proven valuable in RNA-Seq based rare disease studies, due to the rarity of individual phenotypes and the subsequent inability to utilize traditional case vs control group analyses (52). Our work here demonstrates that this utility translates to the study of DNA methylation and has the potential to increase the diagnostic rate of rare disease when utilized alongside existing testing paradigms. DNA and RNA-Seq based analyses in the era of next-generation sequencing have brought about previously impossible genetic diagnoses and it is our hope that expanded sequencing-based methylation testing in this space will represent the next frontier of testing, further lowering the burden of undiagnosed cases.

The methylation profile discovered in Patient 72 represents a potentially novel hypermethylation signature in a rare disease patient. While some enrichment for genes containing features with multiple outlier CpG sites was observed on chromosome chr19 (p-value 3.3^e-3^, OR=1.53) and on chromosomal band chr4p16 (p-value 0.4, OR=2.14), none of the affected genes overlapped with those included in Mayo Clinic epigenetic clinical panel tests, nor did they appear in a collection of known imprinted genes (70). While nascent resources exist to catalog and compare methylation signatures derived by array-based protocols (71), similar resources do not yet exist for sequencing data to our knowledge. Hypermethylation signatures have previously been reported in HMA Syndrome, MRXSN Syndrome and *SETD1B*-related disorder (46) but these are all distinct from the Patient 72 phenotype. Collectively the evidence at hand suggests a likely multigenic effect as the result of increased hypermethylation in hundreds of genes across the genome, rather than results of a focal effect at one or two genes with strong links to the patient phenotype. While this hypermethylation profile might be brought about by a single genetic mutation, we have been unable to identify a causative event. A re-examination of the prior exome sequencing revealed no mutations of interest in known epigenetic genes (72) that might underly a methylation disorder phenotype. Whole genome sequencing has the potential to offer greater insight into potentially causative genomic alterations and this will be pursued as part of the ongoing research-based investigations for Patient 72. Ultimately while the RNA results from OUTRIDER provide an initial validation of results provided by BOREALIS and their functional links to gene expression, the downregulation of these genes in isolation allow us to infer little about the mechanisms creating the patient phenotype.

The current study focused primarily on a single patient as a proof-of-principle, but it must be remembered that widespread, significant results were highlighted for multiple patients in the cohort, and these are undergoing further investigation. Isolated focal changes at the single-site level may also be clinically informative, and the ability to detect these is a strength of BOREALIS and sequencing based methods. However, the burden for follow-up investigation and functional validation quickly becomes unmanageable in the context of a large cohort. Single-site outliers falling within transcriptional control-linked regions of phenotypically relevant genes might logically be of interest, but such findings likely need paired with evidence of aberrant gene expression, which will require routine combination of RNA and methylation assays. Ultimately the results highlighted in this study will be utilized by internal clinical and research teams to inform best next steps and future clinical testing. Patients 36 and 55 for example were identified as outliers based on several analyses and will likely represent priorities when pursuing further testing.

While the analyses described here illustrate a blueprint for methylation analysis from sequencing data in the rare disease field, the data are high-volume and complex and many alternative and potentially valuable approaches likely exist for summarizing and exploring the data. These options will continue to be explored. It is our hope that the present study provides both the computational methodology to make analysis possible, as well as a blueprint for downstream investigation that will continue to be expanded and refined by the research community.

Our work here raises several practical considerations as the field moves toward more routine genome-wide methylation sequencing in rare disease diagnosis efforts. One issue is the interpretation and validation of results derived from approaches like ours. As stated previously, a definitive causative diagnosis for Patient 72 remains elusive despite discovery of a widespread hypermethylation signature. If follow-up whole genome sequencing were to be unsuccessful in resolving the case, the question would remain of what genomic cause, if any, underlies the patient’s phenotype. It is theoretically possible that an epigenetic origin might exist in the absence of a genomic mutation. However, while a single DNA mutation may be sufficient to yield a confirmed clinical diagnosis, the discovery of single or even widespread methylation alterations are often insufficient evidence to do so, due to less thorough understanding and cataloguing of pathogenic events. Carefully planned follow-up functional validation work may be required, creating a financial and logistical burden for patients and institutions. Ultimately it is likely that such challenging cases will persist for years to come but it is our hope that these will be steadily reduced by the routine combination of multiomics testing modalities and the development of public databases cataloguing pathogenic events discovered by different testing paradigms.

Another issue raised by our work is the question of what constitutes the ideal biological sample for testing. In our cohort of 94 patients, we pursued testing of PBMCs in the hope of attaining a homogeneous cell population. Whole blood meanwhile was utilized in RNA-Seq testing due to the necessity to compare to an existing cohort sequenced from whole blood. While the matched whole blood is optimal for comparability with OUTRIDER, it introduces some inconsistency when comparing to methylation data from PBMCs. Furthermore, the use of whole blood for RNA testing limits the number of genes we can successfully profile due to tissue-specific expression, but nonetheless it is readily available from routine clinical blood-draws and has proven utility in the field. As we move toward broader multiomics testing and cohort creation, sample selection will be a problem that will have to be considered and addressed in the context of availability, comparability, and utility. The use of whole blood and PBMCs in our investigation of Patient 72 nevertheless enabled us to demonstrate statistically significant overlap between RNA and methylation results, but further optimization of experimental setup may be possible to maximize the information yielded.

Finally, while our study utilized a relatively large cohort of 94 individuals, our power analysis demonstrates the utility of BOREALIS with smaller numbers of individuals. This characteristic of our approach ensures wide applicability and sufficient power even when studies are limited by small numbers of methylation samples. It is our hope that this will enable wider adoption of our method and be conducive to further discoveries in the field. Ultimately, we hope the ready-availability of our algorithm as a Bioconductor package, and the work described here will serve as a foundation for much future work by ourselves and others.

## Conflict of Interest

The authors declare that the research was conducted in the absence of any commercial or financial relationships that could be construed as a potential conflict of interest.

## Author Contributions

GRO planned the study, performed analysis and interpretation, and wrote the manuscript.

WGJ planned the study, created the software, wrote and reviewed the manuscript.

RJO helped plan the study, selected the patient cohort and reviewed the manuscript.

LSR helped plan the study, selected the patient cohort and reviewed the manuscript.

EWK planned the study, selected the patient cohort and reviewed the manuscript.

## Funding

All funding was provided by the Mayo Clinic Center for Individualized Medicine.

## Data Availability Statement

BOREALIS model files and patient data required to reproduce analysis are provided in Supplementary File 1

## Notes

### Competing Interest Statement

The authors have declared no competing interest.

