## Supplementary figures and images for "Detection of outlier methylation from bisulfite sequencing data with novel Bioconductor package BOREALIS"

### Supplementary Figure 1

#### Slide 1
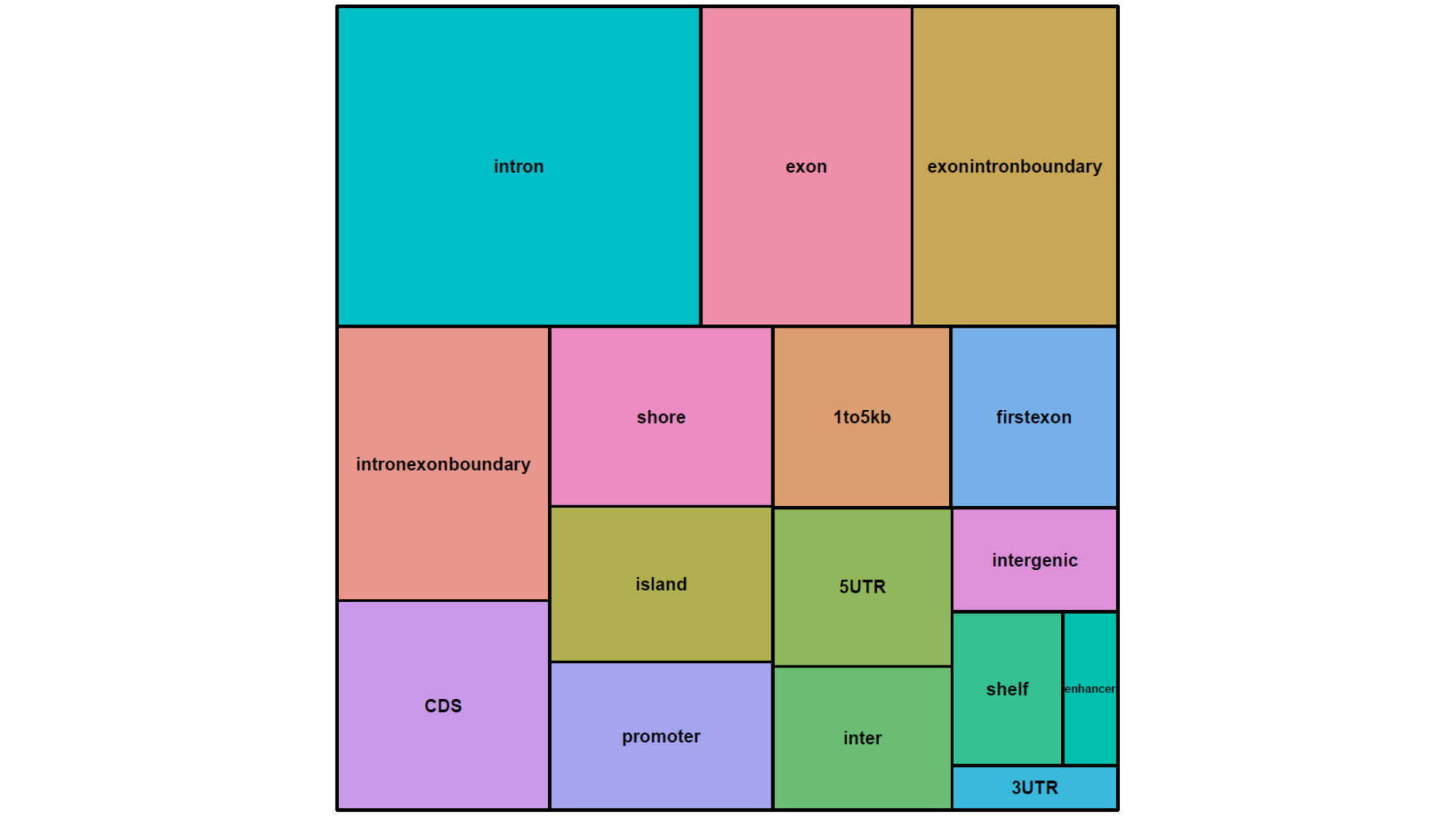
